# Computational constraints underlying the emergence of functional domains in the topological map of Macaque V4

**DOI:** 10.1101/2024.11.30.626117

**Authors:** Dunhan Jiang, Tianye Wang, Shiming Tang, Tai-Sing Lee

**Affiliations:** School of Computer Science and Mellon College of Science, Carnegie Mellon University, Pittsburgh, PA 15213; School of Life Sciences, Peking University, Beijing, China, 100871; Peking-Tsinghua Center for Life Sciences, Beijing, China, 100871; IDG/McGovern Institute for Brain Research, Peking University, Beijing, China, 100871; Key Laboratory of Machine Perception (Ministry of Education), Peking University, Beijing, China, 100871; Computer Science Department and Neuroscience Institute, Carnegie Mellon University, Pittsburgh, PA 15213

**Keywords:** macaque V4, organizational principle, texture and shape, retinotopy, self-organizing map

## Abstract

V4, an intermediate visual area in the ventral visual stream of primates, is known to contain neurons tuned to color, complex local patterns, shape, and texture. Neurons with similar visual attribute preferences are closely positioned on the cortical surface, forming a topological map. Recent studies based on multielectrode arrays and calcium imaging revealed the macaque V4 has neuronal columns tuned to specific natural image features, and these columns are clustered into various functional domains. There are domains tuned to attributes generally associated with object surface properties such as texture or color, as well as domains associated with the shape and form of object boundaries reminiscent of the blobs and inter-blobs in the primary visual cortex. Here, we explored the computational constraints underlying the development of the V4 topological map. We found that the map learned based on self-organizing principles constrained by neuronal column’s tuning and retinotopy position can account for many characteristics of the observed V4 map, including the interwoven organization of texture and shape processing clusters. These anatomical clustering, with the implied local recurrent connectivity, might facilitate a modular parallel processing of surfaces and boundaries of objects along the ventral visual system.

## Introduction

The ventral visual stream extracts a variety of visual attributes to support object recognition (1–3). V4, an intermediate visual area along the ventral stream, is known to contain neurons with selectivity to color (4), orientation (5), spatial frequency (6, 7), curvature (8, 9), figure-ground segregation (10), texture (11–14), and shape information (15– 17). It is found that there are distinct groups of neurons processing texture and shape information rather independently (18, 19). Recent studies further revealed that V4 neurons encode a wide variety of natural image features, and they are organized in columns of distinct functional groups with their tunings ranging from complex patterns such as curvatures to texture and boundary, color, and even facial parts such as eye features (20). Wide-field imaging (21) revealed that the top layer “pixels” of V4 are organized into functional domains.

These pixels are about 90μm x 90μm in resolution, roughly in the scale of a column, and thus each reflecting the feature tuning of neurons in a column, analogous to a V1 orientation column. Some of these feature columns are characterized with distinct surface information, such as texture and color. Other feature columns are embedded with shape information, characterized by object boundaries. These neuronal columns cluster together to form functional domains. The functional domains with surface properties and those with shape properties are interleaved in the map, akin to the blob and inter-blob in the primary visual cortex (21), suggesting there might be distinct modules for parallel surface and shape processing.

What are the logic and constraints underlying the development of the topological map in V4, particularly in the organization of shape and surface processing modules? Along the hierarchical ventral visual pathway, the topological map organization in each area has to resolve the tension that arises from trying to fit multiple visual feature dimensions within a single retinotopic map (22–24). In V1, orientation, spatial frequency, and color feature dimensions are fitted in the spin-wheel hyper-columns of the global retinotopic map. As the neuron receptive field becomes larger going up the visual hierarchical system, the retinotopic constraint becomes looser, and the topological map becomes more dominated by visual features. In the object recognition areas such as the inferotemporal cortex (ITC), the cortical columns become organized in various object-centric functional domains (25) such as animacy (26), object sizes (27), scenes (28), and faces (29–31). While ITC neurons are believed to be organized on the cortical map with ordered features (32–34), retinotopy is still detectable (35–39). V4, an intermediate visual area, has to accommodate the encoding of a much larger set of feature patterns under the confinement of retinotopy (40). The observed topological organization of the functional domains in V4 likely results from the interplay of these constraints. Here, we used a self-organizing algorithm simulated cortical map to better understand these constraints.

The self-organizing map is a computational model based on Hebbian learning, local recurrent excitation, and global inhibition to explain the development of a topological map in the brain (41, 42). It is a process that brings neurons with similar tunings together in a cortical spatial neighborhood. Self-organizing algorithms and models sharing similar assumptions have been used to explain the formation of V1 orientation columns and the ITC object-centric functional domains (43–45). One rationale for this design is to minimize connections (46) between neurons encoding semantically related concepts together for encoding the relationship between these concepts. Thus, topological maps provide a valuable window into neural codes and cortical computation. However, the self-organizing map approach has not been applied to V4 topological maps. A recent spatially regularized topological deep learning network (TDANN) purports to explain cortical map formation throughout the ventral stream from V1 to ITC (45). Yet, the V4 map predicted from this model fails to match some of the key characteristics of the topological map of functional domains we observed in V4, suggesting either some key computational constraints are missing or perhaps there are inadequacies in the algorithm.

Given the measured neuronal columns with different tunings to color natural images, what would be the constraints essential for the self-organizing algorithm to map these neuronal columns to reproduce the observed topology of V4 functional domains (21)? We found that balancing the retinotopic constraint and the feature tuning similarity constraint is key to reproducing a topological map of the functional domains similar to the observed map in many measures, particularly in terms of (1) the distributions of size and adjacency relationship of the functional domains in the map, (2) the simultaneous harboring of a continuous feature map and retinotopic maps, and (3) spatial arrangement of neural clusters with tunings associated with surface properties such as texture and color, and clusters with tunings associated with shape properties such as curvature, junctions and lines. In this paper, we will be referring to this retinotopically constrained self-organizing map as RSOM and a self-organizing map constrained only on image tuning as SOM. Our findings provide further evidence in support of the self-organizing principle as the governing rule in the development of the topological organization of neuronal columns and domains. However, what biological processes underlie the formation of topological maps and how such maps enhance neural computation are mysteries that remain to be unveiled.

## Results

### From the macaque V4 to our RSOM simulated cortical map

To organize all neuronal columns in the V4 digital twin (21) shown in Figure 1A, we leveraged the self-organizing algorithm to train our RSOM mutually constrained by their predicted tuning and estimated retinotopy. With balanced mutual constraints, we allow the Gaussian neighborhood to be big at first and gradually shrink, similar to limiting the wiring cost (46) (see Materials and Methods). This process is biologically plausible as it starts with a long-range horizontal connection that branches to facilitate short-range connections (47, 48). The resulting simulated cortical map in Figure 1E consists of 60 by 60 grids, each learning a tuning curve. We first ask whether the shape of these learned tuning curves is similar to that of the V4. We sort every tuning curve to compute an averaged curve along with a standard deviation from all RSOM grids. This averaged tuning curve responds to 1157 images before dropping to half of the maximum response, while the V4 does so for 842 images out of 50000 images (Figure 1C), corresponding to a sparse code (49). When comparing the averaged tuning of the V4 against our RSOM, it gives a 0.996 Pearson correlation. This drops to 0.992 when comparing the SOM to the V4 benchmark, indicating the retinotopy constraint facilitates the learning of more V4-like tuning curves.

**Fig. 1.**
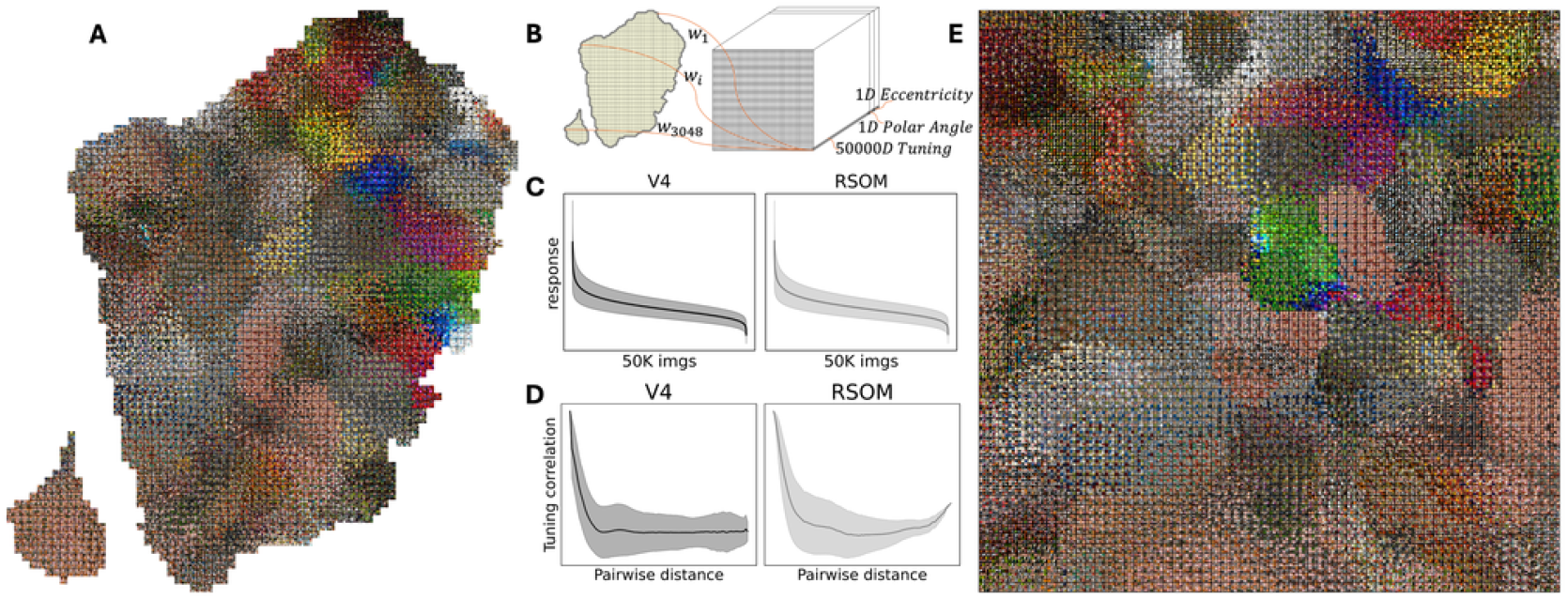
**A**. Monkey V4 digital twin image preference map, each column shows its most preferred nine images out of 50000 color natural images. **B**. RSOM connects to neural column image space. Each column is described by a 50000D tuning curve and its weighted polar angle and eccentricity values. A 60 by 60 gridded simulated RSOM cortical map thus learns a weight vector of 60 by 60 by 50002 dimensions. **C**. Averaged sorted tuning curve shape of the V4 digital twin and our RSOM in darker solid lines, with one standard deviation above and below marked by shadowed areas. **D**. Pairwise V4 columns and RSOM grids tuning correlation as a function of map distance, with function average and standard deviation vectors shown. For every grid, we compute the correlation between its tuning curve and all other grid tuning curves, where correlations are ranked based on the order of pairwise grid Euclidean map distance. **E**. RSOM simulated map image preference. Each grid shows its most preferred nine images from its weight vector first 50000D, representing its tuning curve to the same set of images.

Besides a similar tuning shape, how the tuning curve changes across the cortical surface in these maps is another question. We specifically ask how similar are pairwise grid tuning curves as a function of the physical distance between them on the map. For each RSOM grid, we compute the Pearson correlation between its tuning curve and all other grid tuning curves while recording a pairwise grid distance. This vector of tuning correlations is sorted according to the order of the distance vector. Figure 1D shows the averaged correlation values as a function of distance along with a standard deviation, where spatially adjacent grids have much more similar tuning curves as compared with spatially distant grids. This means spatially closed grids respond particularly to their shared most preferred images, whereas distant grids respond selectively to distinct sets of images. Computing the averaged function vector gives a Pearson correlation of 0.95 between our RSOM and the V4, which drops to 0.89 when comparing the SOM to the V4. This indicates the V4 cortical surface may not be a complete tuning continuum as retinotopy constraint plays a noticeable role in driving a simulated map with a more similar tuning correlation pattern against the benchmark.

### The emergence of V4 domains in the RSOM

Since the V4 map is assembled from sixteen different domains (21), we hypothesize the domain boundary area may experience a sharper tuning curve transition as compared to the tuning transition happening within the domain, thus explaining the above benefit of having an extra constraint other than image tuning. We then look for the emergence of V4 domains by assigning the most similar V4 neuronal column onto every RSOM grid such that the grid can borrow the domain label from the V4 column. This is done by computing correlations between an RSOM grid learned tuning curve and all V4 neuronal columns’ tuning curves such that the column with the highest similarity gets assigned to that grid. We observe the emergence of all V4 domains in our RSOM, as Figure 2A heatmaps demonstrates an example curvature preferring domain enclosed by green line. Two columns of images on the left- and right-most show unique images in the set of top nine most preferred images by all RSOM grids or V4 columns within this domain. Both heatmap colorings indicate averaged response magnitude to these sets of unique images. For the RSOM, we access every boundary grid tuning curve correlation against another nearby grid in a square neighborhood, either within or across the home domain, while recording a pairwise distance. The averaged correlation value as a function of distance is lower for grid pairs across the domain than within the same domain, which is the same for the V4 as well as other domains in both maps. This means tuning transitions more smoothly as a continuum within a domain rather than a more sudden change happening at the domain boundary.

**Fig. 2.**
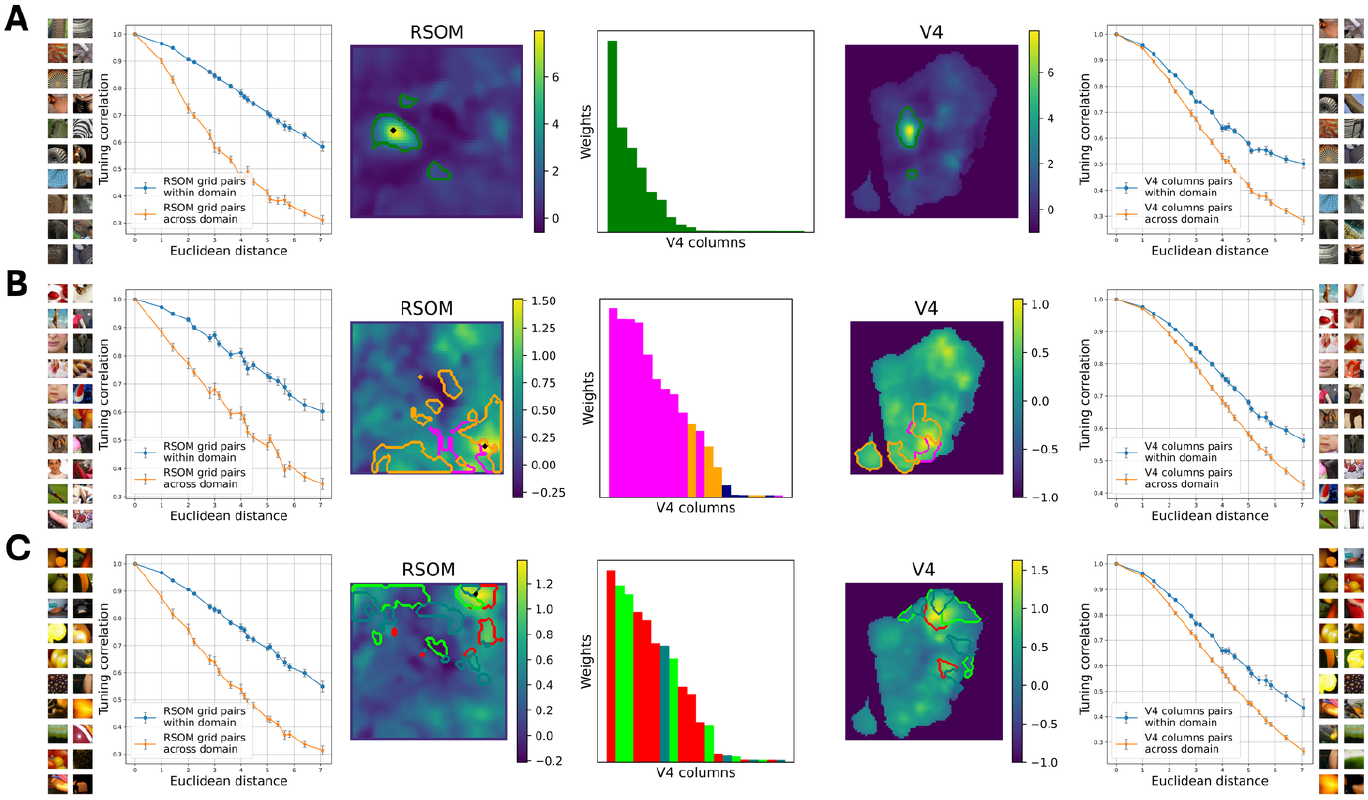
**A**. Texture domain. The left-most images are preferred by RSOM grids as enclosed by green line. They elicit RSOM grid responses, as shown in the left RSOM heatmap. The right-most images are preferred by the same V4 domain with their responses shown in the right V4 heatmap. For all boundary grids, we compute each grid’s tuning correlation to adjacent grids as a function of map distance, the mean and SEM vectors for grids within and across the home domain are shown for the RSOM (left) and the V4 (right). The RSOM example grid’s (a black dot in the heatmap) top one thousand tuning is decomposed as a linear combination of V4 columns through a LASSO linear regression. The middle bar chart shows the top 20 V4 columns with the largest model weights, each as a single bar whose color represents the V4 column domain label. **B**. Face domain. **C**. Curvature domain.

To better understand this perspective, we look at the tuning composition of example RSOM grids at both the center and peripheral of some domains. The black dot in the RSOM heatmap of Figure 2A represents an example grid that has the strongest response inside the curvature-preferring domain, which happens to sit at the domain center. We look at this example grid top one thousand tuning curve and ignore the remaining baseline tuning curve, which is not meaningful. We train an L1-regularized LASSO linear regression model to the fullest to represent its tuning as a linear combination of all V4 column tuning curves. We display the top 20 V4 columns with the biggest linear regression model weights as 20 bars in the center bar chart where its color corresponds to the V4 columns domain label. This example grid is thus learning a tuning curve as a linear combination of only a few V4 columns entirely coming from the same domain in V4.

This story becomes more complicated for an example grid on the more peripheral area, instead of the center, of a shape-preferring domain enclosed by the pink line in Figure 2B heatmaps. While this domain is adjacent to another face-preferring domain enclosed by an orange line, the example black grid in the RSOM heatmap is learning a tuning curve from a slightly more sophisticated subset of V4 columns, including both some home domain columns and a few other columns coming from its adjacent domain. In Figure 2C, we show an extreme case where the example black grid in the red line enclosed domain sits directly adjacent to another two domains that prefer red images (enclosed by the dark green line) and curvature images (enclosed by the light green line). It learns a tuning curve from a very diverse set of V4 columns coming from these three domains. We thereby argue the V4 map is not a perfect tuning continuum, with tuning transitions being smoother within a domain and undergoing more sudden changes at domain boundary lines, as captured by our RSOM. This hints that the V4 map domain assembly and adjacency pattern should not be completely driven by the image tuning constraint.

### Simulated cortical map domain adjacency, retinotopy, and feature dispersity patterns

We visualize domain label maps given by the V4 column assignment in Figure 3A, where each color indicates a specific domain that can split into a few components across the map. All domain component sizes are visualized as a bar chart in Figure 3B with the same coloring. The average domain component sizes of the RSOM and V4 achieve a Pearson correlation of 0.791, whereas this correlation drops to 0.427 for the SOM. We use a square domain adjacency matrix to characterize cortical map domain assembly pattern, where the *i*th row and *j*th column value indicates the number of grids in the *i*th domain being adjacent to grids in the *j*th domain, where each row is normalized. The Pearson correlation between the flattened adjacency matrix of the RSOM and the V4 reaches 0.755, whereas this value drops to 0.484 for the SOM. These findings indicate the crucial role of retinotopy in driving the simulated cortical map to be more V4-like, where it guides domains into their relative positioning onto the cortical map. This perspective matches evidence in the ITC pointing to an experience-insensitive and predetermined innate network structure (50, 51), although similar domain adjacency patterns were only studied in the ventral visual pathway at a macro-scale (52, 53).

**Fig. 3.**
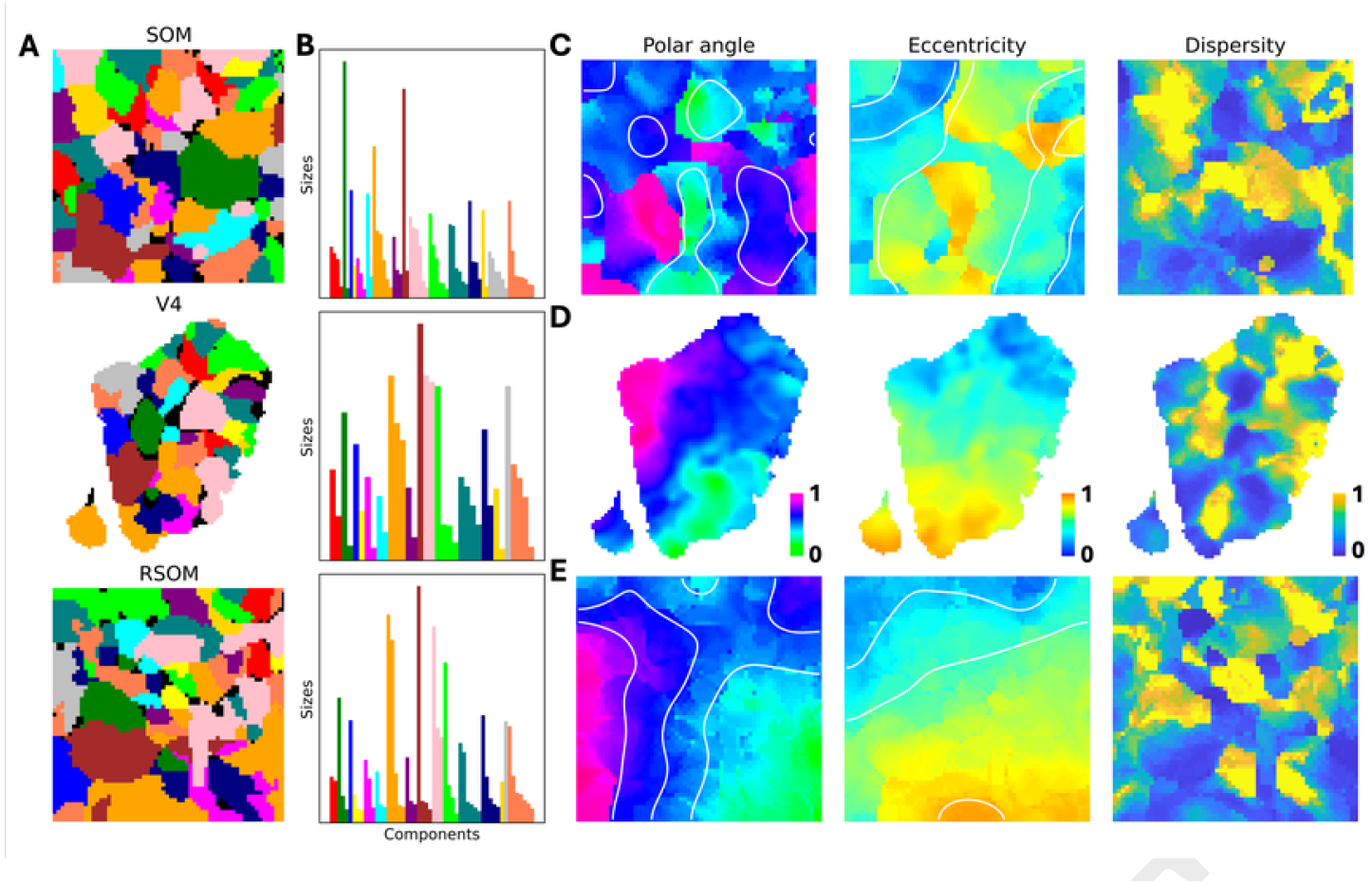
**A**. Domain label map by assigning V4 column onto SOM/RSOM grids. Sixteen distinct colors each represent one domain. Black region means erased grids/columns, whose original connected-component with the same domain label is not large enough. **B**. Domain connected-component sizes. Each bar represents a single component where the same set of colors are used across subfigures to indicate domain label. **C**. SOM polar angle, eccentricity, and feature dispersity maps from V4 column assignment, with contour lines added for the Gaussian smoothed retinotopy maps, indicating their global pattern. **D**. V4 retinotopy and dispersity maps. **E**. RSOM retinotopy and dispersity maps, with contour lines similarly added.

V4 column assignment also brings retinotopy and feature dispersity measurements for every simulated map grid. We visualize these map patterns for the SOM (Figure 3C), V4 benchmark (Figure 3D), and our RSOM (Figure 3E). Due to the retinotopy constraint, contour lines in the polar angle map intersect with those lines in the eccentricity map roughly perpendicularly in the RSOM simulated cortical map, similar to the V4, instead of the chaotic pattern found in the regular SOM. For the dispersity pattern, we did a two-way hierarchical clustering on all maps to cluster grids into either low or high dispersity types. This finds the connected high dispersity components in all maps, and their sizing statistics are shown in Table 1. Although there is no explicit dispersity driving force in our RSOM, the feature dispersity map splits into many high dispersity components in yellow that prefer dispersed texture features and low dispersity areas in blue that tune to more spatially localized shape and object boundaries, resembling the high- and low-dispersity segregation in V4. Without the retinotopy constraint, the regular SOM features less segregated high dispersity components, where one large high dispersity cluster emerges across the map instead. This exciting finding points to a retinotopy-dependent organizational principle being the driving force behind the observed modular parallel processing of segregated texture and shape clusters in the V4 and along the ventral visual processing stream.

**Table 1.** High dispersity component size statistics, including the total number of components, total map area ratio, averaged component area ratio, standard deviation of all component area ratios, minimum component area ratio, and maximum component area ratio.

| Cortical Sheets | num | sum | mean | std | min | max |
| --- | --- | --- | --- | --- | --- | --- |
| 1. V4 | 7 | 0.31 | 0.04 | 0.04 | 0.01 | 0.09 |
| 2. SOM | 6 | 0.32 | 0.05 | 0.06 | 0.01 | 0.17 |
| 3. RSOM | 9 | 0.33 | 0.04 | 0.03 | 0.01 | 0.09 |

### TDANN purported V4 and ITC layer map topologies

As supervised models tend to encode images differently from the brain (54, 55), innovation of spatially regularized goal-driven network (45) was trained from contrastive learning to model the ventral visual stream cortical map. Here, we examine TDANN’s simulated V4 and ITC cortical maps. For each map, we extract all artificial unit responses to the same set of 50000 natural images. The simulated cortical map area was segmented into 60 by 60 pixels such that each pixel have an aggregated tuning curve from all artificial units within this pixel area. We show the top nine most preferred images of these pixels for both the V4 (Figure 4A) and the ITC (Figure 4B) layer. Upon an initial examination, the V4 map forms small domains tuned to color and object but no faces, while the ITC map forms larger domains only at the map center tuned to color, face, and object. While this TDANN model was not trained to match the macaque V4 neural code, we examine its tuning curve shapes. We found its ITC average tuning curve shape achieves a 0.947 correlation against our benchmark macaque V4, which drops to only 0.708 for its purported V4 layer, whose tuning curve also becomes much less sparse. We repeat the pairwise pixel tuning curve correlation as a function of simulated cortical map distance analysis and find 0.838 and 0.787 correlations for the purported V4 and ITC layers against our benchmark V4 (Figure 4C and 4D). Due to distinct image preferences, we select all unique top nine most preferred images from macaque V4 neuronal columns, to build a representation similarity matrix for every cortical map. This gives a correlation of 0.303 for the TDANN V4 map and a 0.242 correlation for the TDANN ITC map against the benchmark V4, indicating a divergence between goal-driven model image representation and the biological visual cortex representation on a population code level (56).

**Fig. 4.**
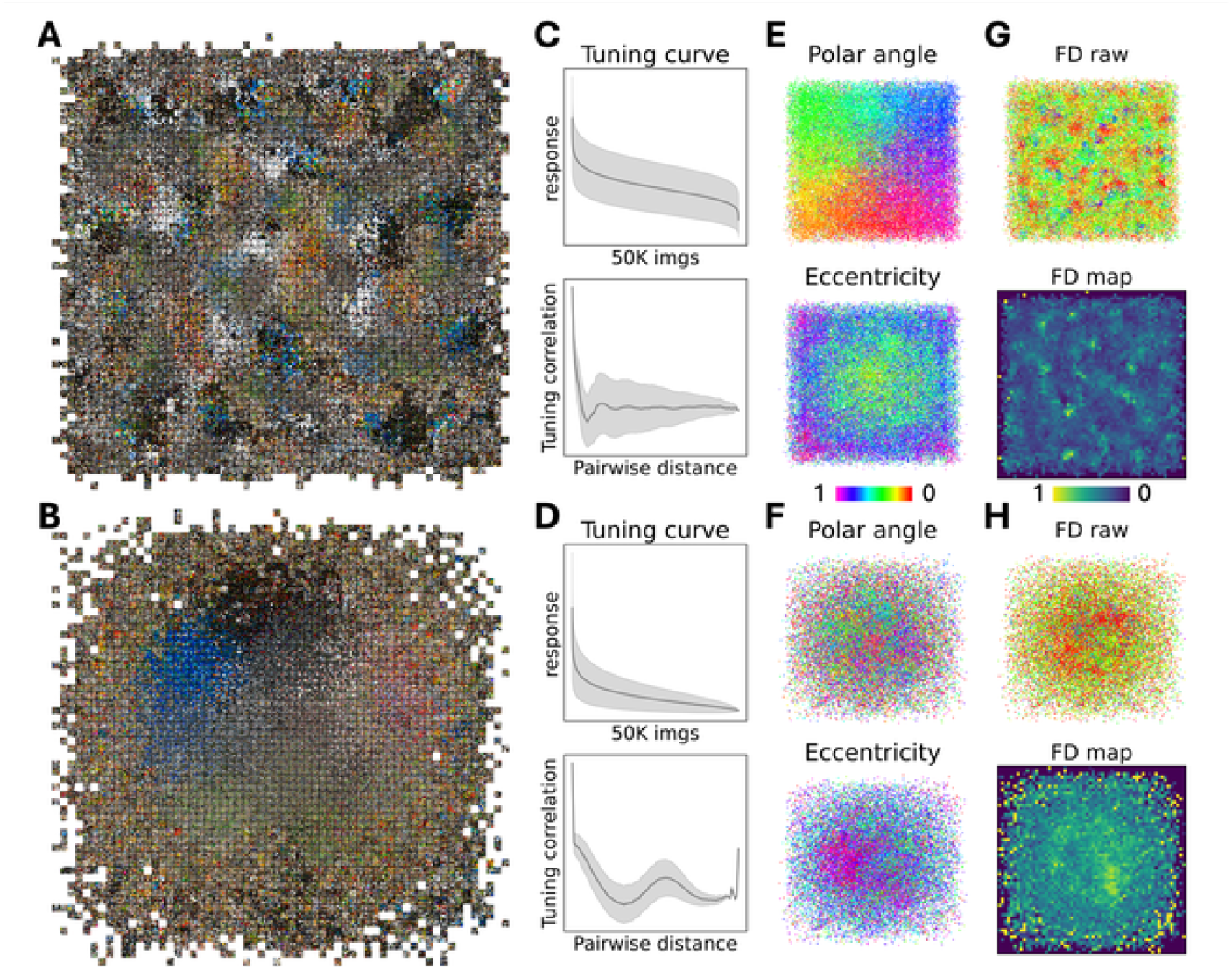
**A**. TDANN purported V4 image preference map, with 60 by 60 pixels each showing nine most preferred images. This cortical map was equally segmented into pixels, where all units within a pixel yield a mean tuning. **B**. TDANN purported ITC 60 by 60 image preference map. **C**. Top: averaged sorted tuning curve for V4 layer pixels with a much wider shape than that of the macaque V4 (corr=0.708). Bottom: V4 layer pixels pairwise tuning curve correlation as a function of map Euclidean distance (corr=0.838 against the macaque V4). **D**. Top: ITC layer pixels mean tuning has a similar shape against the macaque V4 (corr=0.947). Bottom: ITC layer pixels pairwise tuning correlation as a function of distance (corr=0.778). **E**. Top: V4 layer polar angle map. Bottom: V4 layer eccentricity map. **F**. Top: ITC layer polar angle map. Bottom: ITC layer eccentricity map. **G**. Top: V4 layer feature dispersity map. Bottom: aggregated V4 layer feature dispersity map with 60 by 60 pixels. **H**. Top: ITC layer feature dispersity map. Bottom: aggregated ITC layer feature dispersity map with 60 by 60 pixels.

We further tested the retinotopy organization in these simulated cortical maps, as the position of artificial unit was retinotopically initialized based on their feature map location in the convolution structure. Here, we use this convolution layer feature map structure to assign retinotopy positions for all units based on their position in the feature map, assuming the center of this feature map is the fovea of the visual field. We show both polar angle and eccentricity values for all artificial units in the purported V4 (Figure 4E) and ITC (Figure 4F) layers, indicating some retinotopy preserved at the V4 simulated map but not at the ITC map where long-range horizontal connection and neighborhood position optimization likely disrupted the retinotopic structure. But whether the preserved retinotopic organization at the simulated V4 cortical map gives rise to segregated shape and texture parallel and modular processing remains unknown. We applied the same feature attribution analysis (21) to investigate the feature dispersity properties of artificial units in both the V4 and the ITC layers. As a result of our test images, where the peripheral portion was occluded to mimic what the monkey saw during the experiment, we chose the center feature map artificial unit as a representative of all other units in the same feature channel, disregarding their feature map locations. However, starting from the purported V4 layer to the ITC layer, we found TDANN units’ feature dispersity values follow a different distribution from the monkey V4 neuronal columns in the digital twin. As a result, we did not observe the high and low feature dispersity segregation across both simulated cortical maps here in Figure 4G and 4H, indicating different feature processing and learning between the TDANN and the macaque V4.

## Conclusions

In this paper, we showed that the recently observed V4 topological map of natural image feature selectivity is compatible with the self-organizing algorithm advanced by Kohnon (41). Self-organizing and similar algorithms have been used to explain V1 and ITC maps (44, 57, 58), which have not been used to explain V4 maps, particularly the topological natural image preference map (21). Using a digital twin built by fitting wide-field calcium imaging neural responses over a large stretch of V4 at 90 μm scale to a large set of natural images, we have come to a better understanding of how natural scenes are embedded in V4 in the form of a topological map. Here, we show that the self-organizing algorithm with retinotopic constraint, as well as feature similarity constraint, is sufficient to reproduce cortical maps that match the observed V4 map of functional domains of natural image feature preferences in a number of metrics, particularly in terms of (1) the distributions of size and adjacency relationship of the functional domains in the map, (2) the simultaneous harboring of a continuous feature map and a continuous retinotopic map, and (3) spatial arrangement of neural clusters with tunings associated with surface properties such as texture and color, and clusters with tunings associated with boundary shape properties such as curvature, junctions, and lines.

While the functional roles of the observed cortical topology and the clustering of domains remain uncertain, minimizing horizontal connection (46) and top-down feedback connections to encode related concepts to facilitate neural computation is likely a key factor. Visual independent factors such as other sensory modalities and memories that affect the functions of the visual cortex might also contribute to the formation of topological maps, as there are long-range connectivity extending outside of the visual cortex (59–61) and functional domains in extrastriate visual cortex can be activated by concepts communicated with sound in the congenitally blind group (62). Retinotopy, as a visual dependent factor, is known to be an important constraint guiding the formation of the visual cortical map. Current evidence suggests the existence of retinotopy organization of the visual space in primate visual systems at birth, even prior to the formation of neuronal selectivity and functional domains (22–24). Traces of such inborn retinotopy organization are present at matured primate V4 (40). Eccentricity remains an important factor (35–39) in cortical maps of the ITC, where even scene domain (28) and face domains (30, 31) within the ITC are found to exhibit retinotopic organization at the mesoscale.

We argue that the organization of multiple visual feature dimensions in the macaque V4 area should also simultaneously satisfy the retinotopic organization projected from the retina visual input. Without this retinotopic constraint, as shown in our SOM, functional domains tend to appear in larger chunks. The pressure due to the retinotopic constraint, as in our RSOM, could break a functional domain, such as the facial feature domain, into multiple pieces, conceptually similar to orientation columns in the different hypercolumns in V1. The retinotopy constraint also breaks the high-dispersity domain clusters into pieces, surrounded by low-dispersity domains. It is possible that the high dispersity domains might derive their major input from the V1 blobs / V2 thin stripes and the low dispersity domains from the inter-blobs / V2 thick stripes, constituting parallel streams of specialized surface (color, texture) and shape (orientation, curvature, and form) processing in the ventral visual system. However, further anatomical and physiological studies are needed to evaluate this conjecture.

On the other hand, neural networks have been shown to match the ventral visual systems reasonably well by measures of neural response prediction and representational similarity analysis (54, 55). Neural response prediction suggests different network layers have a rough correspondence with the different visual areas of the ventral stream in terms of the embedding features, while representational similarities suggest a matching in the geometry of the neural manifold of these embedded features. Deep convolutional neural networks, however, do not exhibit topological maps, which is a key characteristic of the organization of the visual cortex. A recently proposed spatially regularized topological deep artificial neural network (45) is an attempt to address this gap by imposing a loss that encourages neurons with similar tunings to come together spatially but subject to the normal convolutional neural network model architecture. Unfortunately, this model fails to predict the V4 map of natural image features that we observed, though it generates a V1-like map in the early layer and an ITC-like map in the higher layer. The features in the purported V4 layer, specifically block 3.1 in the ResNet18 model, appear to be missing some of the V4 feature tunings, most notably facial feature tunings, as well as a greater variety of color tunings. It does exhibit functional domains but with a coarser retinotopic map and completely scattered and chaotic high and low dispersity clusters, similar to our SOM simulated map with only the tuning feature constraint. This suggests the segregation of high and low dispersity clusters, presumably corresponding to surface and shape processing, requires interaction with a relatively high retinotopic constraint during the learning of the map.

Both self-organizing maps and topological CNN primarily address the issues of understanding the topological map at an algorithmic and representational level (63). The theoretical level account and the implementation level account remain to be explored and developed. At the theoretical level, the topological map might be an efficient embedding of the efficient codes of natural scenes related to the sparse manifold transform (SMT) (64), where neurons encoding spatially and/or temporally co-occurring visual events become more connected due to Hebbian learning and slow-feature analysis (65). The excitatory mutual connection might bring the neurons with close semantic relationships closer together in cortical space to facilitate building connections. At the implementation level, further empirical experiments are required to understand the exact biological processes and mechanisms that realize map formation under the retinotopic and feature similarity constraints.

## Materials and methods

### RSOM parameters and data

We applied a PyTorch-based Kohonen self-organizing algorithm to the image space of V4 digital twin columns, each coupled with its retinotopy, including one polar angle and one eccentricity value. RSOM learns a simulated cortical map of 60 x 60 (H x W) grids where each grid learns a tuning curve. The weight of this RSOM is of shape H x W x F (60 x 60 x 50002), of which the third axis first 50000D learns the tuning and the last 2D corresponds to polar angle and eccentricity. This design allows a direct constraint in finding every best matching unit from both tuning and retinotopy. The training data is 3048 monkey V4 digital twin column vectors, each consisting of 50000D tuning plus its polar angle and eccentricity values.

### RSOM training

We do a complete random weight initialization while normalizing the weight vector for each grid. This ensures no V4 topology information was pre-planted into the simulated cortical map. The RSOM only sees one V4 column vector at a time when training. The minimum Euclidean distance between the input vector and all RSOM grid weight vectors finds the best matching unit (BMU, see Equation 1), where *w* = 1 *if ∈* [1 : 50000] and *w* = 125 *if ∈* [50001 : 50002].

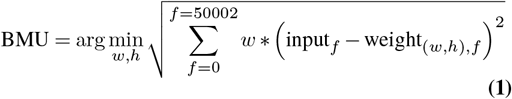

This winning grid serves as the center of a Gaussian neighborhood function in Equation 2 to update all grid weights, with grids closer to the BMU getting more weight changes toward the input V4 column vector while more distant grids get fewer updates. This filter function *η*_*t*_ starts big and decays to an intermediate value. The weight update (Equation 3) uses a constant learning rate *α* on top of the neighborhood function. This RSOM is trained for 120 epochs, where a single epoch iterates through all 3048 V4 neuronal columns for one time. The SOM is similarly trained as well.

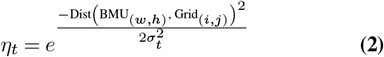

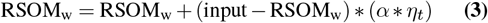

### RSOM example grid tuning composition

For an example grid, we select its top one thousand tuning curve portion. We train a LASSO linear regression model to fit this partial tuning curve as a linear combination of 3048 V4 columns’ partial tuning to exactly these one thousand images (Equation 4). The loss is a mean squared error plus another L1 punishment term applied to all 3048 weights. This regularization is multiplied by 0.01 to control the penalty. Each model is trained with a constant learning rate *α*=0.05 for 7500 epochs to the fullest, giving a final loss around 0.01.

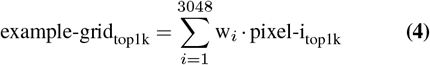

### Map connected components and adjacency matrix

In both simulated cortical maps and the V4 map, we find domains as connected components. This means a cluster of grids that are connected also share the same domain label. We define an eight-connectivity where both direct adjacent and diagonal adjacent grids of the same label will be considered as connected. To eliminate small and noisy components in the map, we filter out those connected components whose size is smaller than ten by erasing the label of these grids. We compute an adjacency matrix for each map to quantify relative domain positioning patterns. This is a 16 by 16 matrix where each row and column correspond to a specific domain. For the *i*th domain represented by the *i*th row, we look at all of its connected components. We survey each component’s adjacent grids where whenever it is adjacent to one grid labeled as the *j*th domain, we add a value of one to the *i*th row *j*th column and the *j*th row *i*th column. Each row vector is normalized after surveying the entire map.

### Code

Scripts for model training and simulated map analysis are available at: https://github.com/777dunhan/Topological-map-of-Macaque-V4

